# *De novo* discovery of traits co-occurring with chronic obstructive pulmonary disease

**DOI:** 10.1101/2022.07.20.500731

**Authors:** E. Golovina, T. Fadason, R.K. Jaros, H. Kumar, J. John, K. Burrowes, M. Tawhai, J.M. O’Sullivan

## Abstract

Epidemiological research indicates that chronic obstructive pulmonary disease (COPD) is a heterogeneous group of chronic lung conditions that are typically accompanied by cardiovascular disease, depression, lung cancer and other conditions. Genome-wide association studies (GWAS) have identified single-nucleotide polymorphisms (SNPs) associated with COPD and the co-occuring conditions, suggesting common biological mechanisms underlying COPD and these co-occuring conditions. To identify them, we have integrated information across different biological levels (i.e. genetic variants, lung-specific 3D genome structure, gene expression and protein-protein interactions) to build lung-specific gene regulatory and protein-protein interaction networks. We have queried these networks using disease-associated SNPs for COPD, unipolar depression and coronary artery disease. Our results show that COPD-associated SNPs can control genes involved in the regulation of lung or pulmonary function, asthma, brain region volumes, cortical surface area, depressed affect, neuroticism, Parkinson’s disease, white matter microstructure and smoking behaviour. We describe the regulatory connections, genes and biochemical pathways that underly these co-occuring trait-SNP-gene associations. Collectively, our findings provide new avenues for the investigation of the underlying biology and diverse clinical presentations of COPD. In so doing, we identify a collection of genetic variants and genes that may aid COPD patient stratification and treatment.

## Introduction

Chronic obstructive pulmonary disease (COPD) is a heterogeneous group of chronic lung conditions that are characterized by persistent respiratory symptoms and airflow limitation due to airway and/or alveolar abnormalities (Global Initiative for Chronic Obstructive Obstructive Lung Disease. 2020). These abnormalities are caused by a combination of distinct pathophysiological processes that result in diverse clinical presentations, responses to treatment, and patterns of progression. According to the World Health Organisation, COPD accounted for more than 3.23 million deaths in 2019 and remains the third leading cause of death worldwide (Global Initiative for Chronic Obstructive Obstructive Lung Disease. 2020).

Given the widespread exposure to the environmental factors (e.g. smoking, indoor and outdoor air pollution, childhood respiratory infections) that contribute to the development of COPD, it is striking that most individuals will never develop COPD. The variance in individual susceptibility to COPD can be partly explained by genetic factors. The estimated genetic heritability of COPD ranges from 20% to 40% for airflow limitation (Stolz 2020; Gim et al. 2020) and up to 60% for smokers (Zhou et al. 2013). This variation in heritability suggests that the presence of two or more conditions (hereafter defined as co-occuring) can increase the risk of mortality in COPD patients.

A better understanding of COPD co-occuring conditions is essential to enable effective management, therapeutic optimization and reduce the costs of managing COPD patients (Mannino et al. 2015). Epidemiological and genetic studies have reported that beyond respiratory impairment COPD-associated co-occuring conditions include coronary artery disease (CAD), lung cancer, osteoporosis, mental health problems such as anxiety, unipolar depression (UD), Alzheimer’s disease (AD), and Parkinson’s disease (PD) (Burke and Wilkinson 2021; Carmona-Pírez et al. 2021; Xia et al. 2020; Martucci et al. 2021; Ställberg et al. 2018; Li et al. 2015; Cavaillès et al. 2013). The presence of these conditions in COPD patients indicates that common or interacting biological mechanisms underlie these conditions.

To date, genome-wide association studies (GWASs) have identified common single nucleotide polymorphisms (SNPs) that are associated with COPD, or its individual co-occuring conditions (Kim et al. 2021; Zhu et al. 2019). The majority of the COPD-associated SNPs are located within the non-coding genome. Therefore, the impacts that these SNPs have on the biological pathways and processes underlying the development of COPD remain unclear. It is possible that the COPD-associated SNPs mark regulatory regions (i.e. expression quantitative trait loci [eQTLs]) that are associated with tissue-specific gene expression. eQTLs can interact with their target genes in three dimensions, forming spatial eQTL-gene regulatory connections that span the genome (e.g. cis, ≤1Mb on the same chromosome; trans-intrachromosomal, >1Mb on the same chromosome; or trans-interchromosomal, between different chromosomes). These spatial interactions are cell and tissue type-specific (GTEx Consortium 2020; Halow et al. 2021). As lung is the primary affected tissue in COPD, integrating lung-specific spatial chromatin interactions and eQTL information may help us understand how SNPs impact biological pathways that increase an individual’s risk of developing COPD.

Little is known about functional relationships between genes and phenotypes in the lung. However, gene regulation is widely understood to occur through the combinatorial action of regulatory elements, transcription factors and genes within complex networks (i.e. gene regulatory network, GRN) (Buenrostro et al. 2018; Chen et al. 2021a; Zaied et al. 2022; Chen et al. 2021b). Moreover, genes encode proteins, that physically interact with each other to form a complex protein-protein interaction network (PPIN) that responds to biological and environmental signals. Here, we integrated COPD-associated SNPs with: 1) information on the genome organization within the lung; and 2) lung-specific eQTL information to identify genes that are spatially regulated within the lung tissue. We integrated information across a lung-specific GRN and PPIN to identify conditions that were co-occuring with COPD. Collectively, our results highlight potential regulatory mechanisms and pathways important for COPD etiology. These results open a new avenue towards understanding the diverse clinical presentations of COPD and patient stratification.

## Results

### COPD-associated SNPs mark putative regulatory regions in the lung

COPD-associated SNPs (*p* < 5×10^−8^, n=263) were downloaded from the GWAS Catalog (Supplemental Tables S1 and S2) and run through the CoDeS3D pipeline (Fig. 1A). Approximately 96% of the identified eQTLs were located within non-coding genomic regions, with 66.02% and 18.45% of them being intronic and intergenic, respectively (Supplemental Fig. S1A and S1B, Supplemental Table S2). Analysis of these SNPs using the CoDeS3D (Fadason et al. 2018) pipeline (Fig. 1A) identified 103 eQTLs and 107 genes that are involved in 151 significant (FDR < 0.01) eQTL-gene interactions within the lung (Supplemental Fig S1, Supplemental Table 3S). The majority of COPD-associated eQTLs (n=67) are involved in one-to-one, 26 eQTLs – in one-to-two and 8 eQTLs – in one-to-three eQTL-gene regulatory interactions (Supplemental Fig. S1C, Supplemental Table S3). Only 2 eQTLs (i.e. rs2277027 and rs9435731) were associated with the regulation of ≥4 genes (i.e. *ADAM19, CTB-109A12*.*1, CTB-47B11*.*3, CYFIP2* and *ATP13A2, CROCC, MFAP2, RP1-37C10*.*3*, respectively; Supplemental Fig. S1C, Supplemental Table S3). The majority of the identified eQTL-gene regulatory interactions (n=148) were cis-acting (Supplemental Fig. S1D, Supplemental Table S3). One trans-intrachromosomal (i.e. rs2077224-*NAV2*) and two trans-interchromosomal (i.e. rs12894780-*LIPC* and rs2128739-*KALRN*) eQTL-gene interactions were identified within the lung (Supplemental Fig. S1D, Supplemental Table S3). Collectively, COPD-associated eQTLs are associated with changes in transcription levels of 84 protein-coding genes, 22 non-coding RNA genes and one pseudogene (Supplemental Fig. S1E).

**Figure 1.**
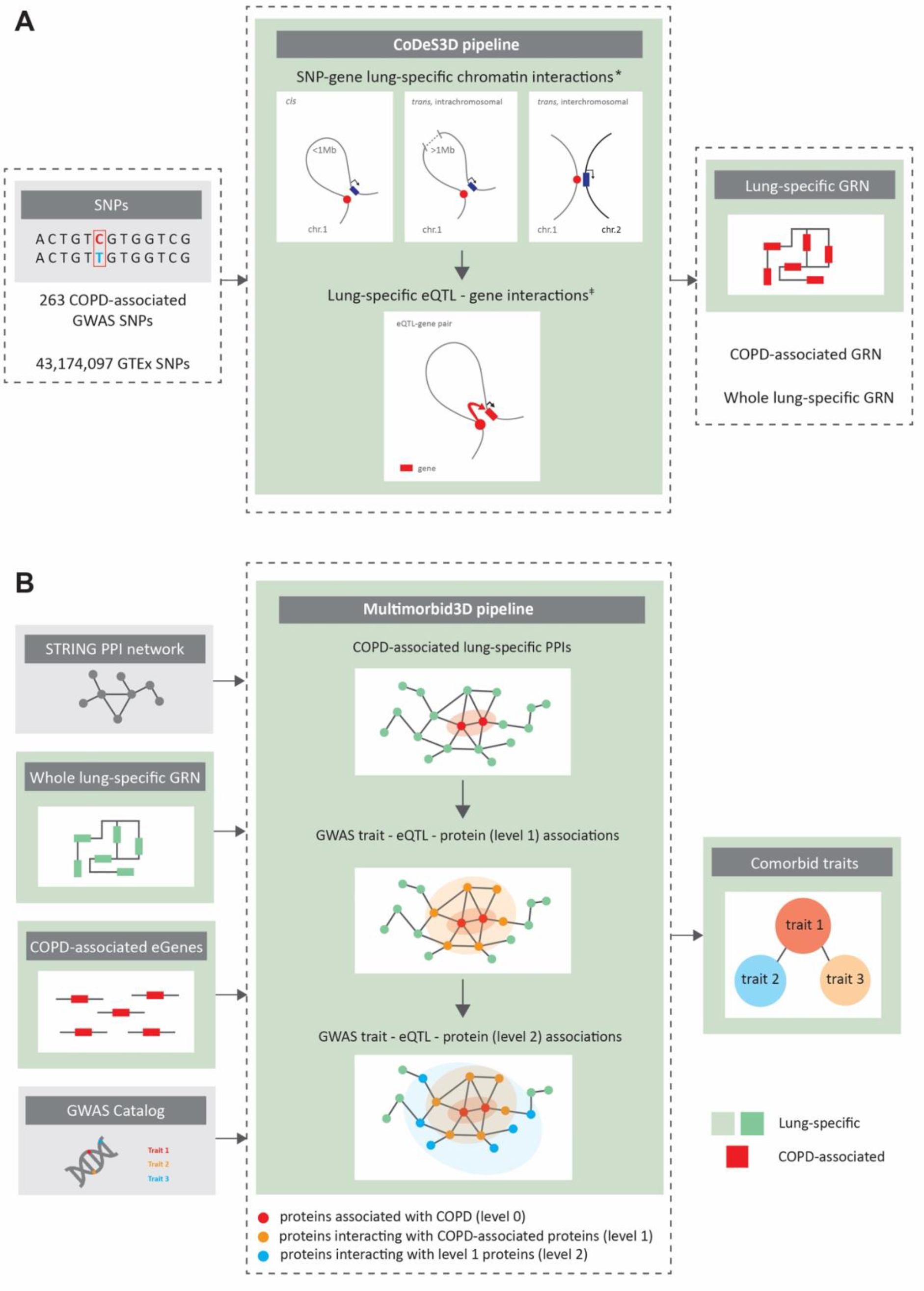
Overview of the analytical pipelines used in this study. (**A)** The CoDeS3D pipeline was used to identify lung-specific gene regulatory networks (GRNs). First, 263 GWAS SNPs associated (*p* < 5×10^−8^) with COPD were run through the CoDeS3D pipeline to identify 151 spatial eQTL-gene regulatory interactions in the lung (COPD-associated lung-specific GRN). Next, all GTEx SNPs (MAF ≥ 0.05, n=43,174,097) were downloaded from dbGaP (Supplemental Table S1) and analysed using CoDeS3D to identify ‘all’ significant lung-specific spatial eQTL-gene regulatory interactions (whole lung-specific GRN). The resultant whole lung-specific GRN is comprised of 873,133 spatially constrained regulatory interactions involving 740,028 eQTLs and 15,855 genes (Supplemental Fig. S2 and S3). (**B)** The Multimorbid3D pipeline was used to identify potential co-occuring conditions associated with COPD. * Hi-C datasets for primary lung cells were obtained from Schmitt et al. (GEO accessions: GSM2322544 and GSM2322545). ǂ eQTL datasets for lung was obtained from GTEx v8 (dbGaP accession: phs000424.v8.p2).

### COPD-associated genes are enriched for diverse biological processes in the lung

Functional gene ontology enrichment analysis identified metabolic, behavioural, regulatory and protein modification processes (e.g. “phosphorus metabolic process”, “behavioral response to nicotine”, “regulation of postsynaptic membrane potential”, “protein acetylation” and “protein acylation”) as being significantly (*p* < 0.05) enriched within the 107 COPD-associated genes (Supplemental Table S4, Supplemental Fig. S4). These 107 genes encoded proteins that formed 9 COPD-associated lung-specific protein-protein interaction clusters (Fig. 2, Supplemental Table S5). Louvain clustering identified nine highly connected modules within these 9 clusters (Fig. 2). Pathway analysis of the nine highly connected modules identified seven that were enriched (FDR < 0.05) for immune pathways, sulfur relay system and folate biosynthesis, axon guidance and regulation of actin cytoskeleton, focal adhesion and regulation of actin cytoskeleton, insulin resistance and insulin signaling pathway, TGF-beta signaling pathway, neuroactive ligand-receptor interaction and cholinergic synapse. Two modules were of unknown function (Fig. 2).

**Figure 2.**
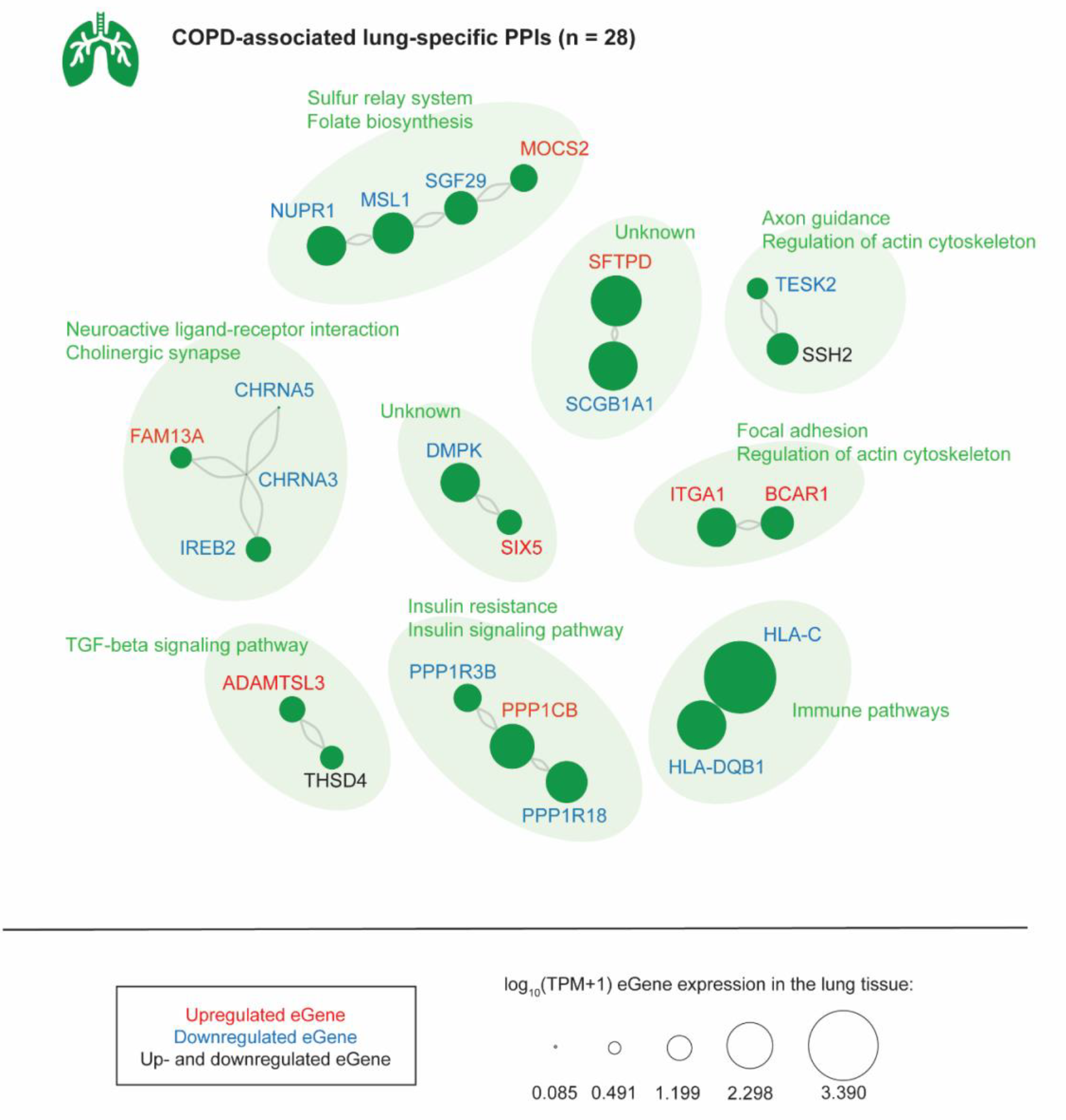
Louvain clustering identified nine PPIN modules within the lung: seven were enriched for biological pathways, and two modules with unknown pathway enrichment. Significantly enriched (FDR ≤ 0.05) pathways included: immune pathways, sulfur relay system and folate biosynthesis, axon guidance and regulation of actin cytoskeleton, focal adhesion and regulation of actin cytoskeleton, insulin resistance and insulin signaling pathway, TGF-beta signaling pathway, neuroactive ligand-receptor interaction and cholinergic synapse. Red text, the eQTLs is associated with upregulation of the gene; blue text, the eQTL is associated with downregulation of the gene transcript; black text, eQTLs are associated with up- and downregulation of the gene transcript levels.

### COPD has associations with co-occuring traits in the lung, brain and blood

Patients with COPD often also suffer from cardiovascular disease, osteoporosis, lung cancer, sleep disorders and mental health problems (Burke and Wilkinson 2021; Carmona-Pírez et al. 2021). Yet the biology of these interactions is rarely known. The Multimorbid3D algorithm was used to integrate COPD-associated genes, the LSPPIN, the lung-specific GRN and all catalogued GWAS SNP-trait associations (30/03/2022) to identify co-occuring conditions and potential regulatory mechanisms underlying these associations with COPD (Fig. 1B). We identified 39 GWAS traits that are significantly (FDR ≤ 0.05) enriched for eQTLs that target the COPD-eQTL associated genes (“level 0”; Fig. 3, Supplemental Table S6). Most of the level 0 co-occuring traits were “lung-” (i.e. COPD, lung function, pulmonary function, post bronchodilator FEV1, asthma), or “mood/brain-related” (i.e. brain region volumes, cortical surface area, depressed affect, neuroticism, Parkinson’s disease, white matter microstructure, smoking behaviour). eQTLs that regulate genes encoding proteins within levels 1-4 of the expanded COPD lung protein interaction network were enriched within traits that have and have not been previously recognized as being co-occuring with COPD (Fig. 3).

**Figure 3.**
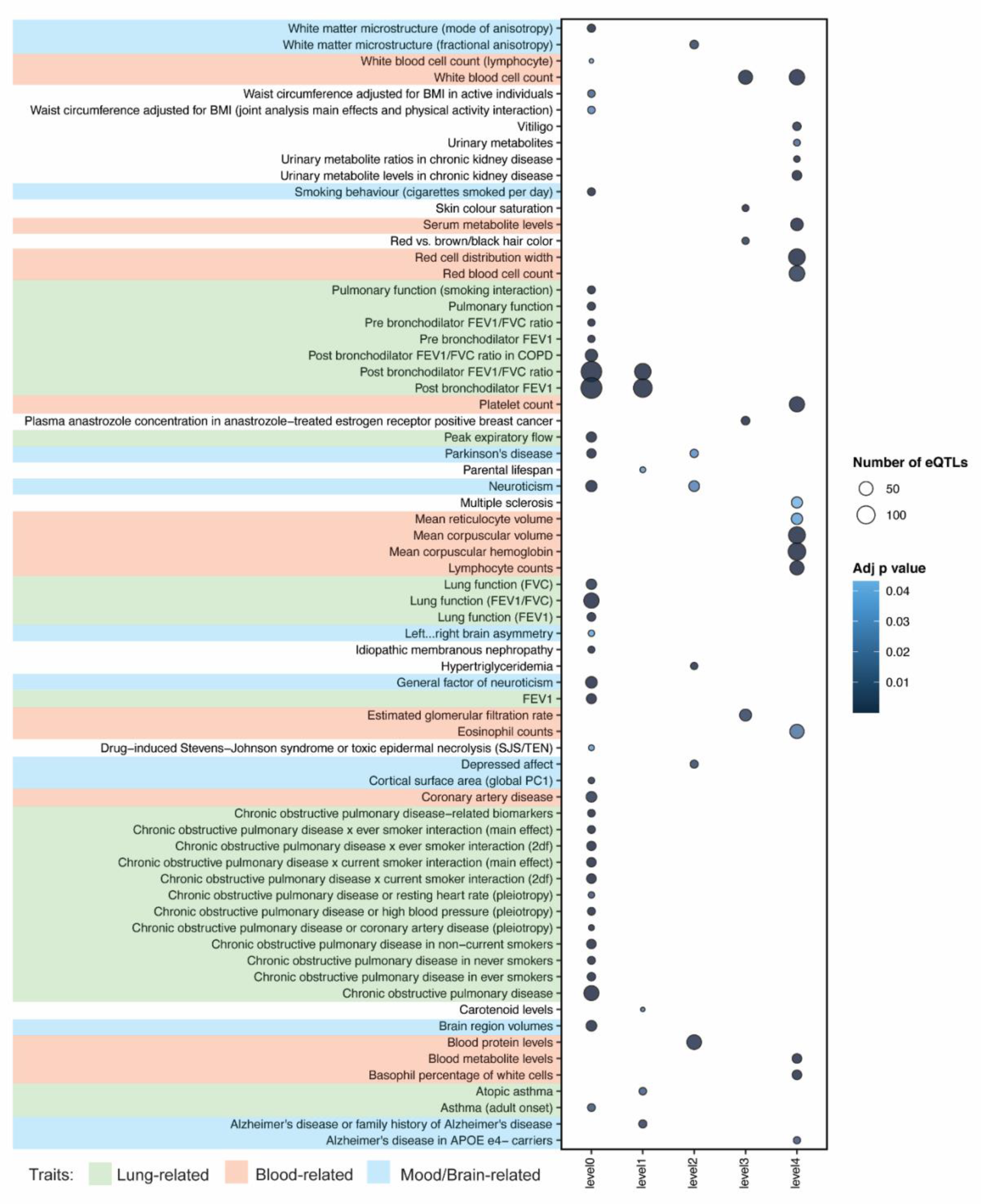
Network analysis identified co-occuring conditions that are associated with COPD. We identified 39 GWAS traits that are enriched (FDR ≤ 0.05) for eQTLs associated with COPD-eQTL target genes (level 0). Most of these co-occuring traits are “lung-related” (i.e. COPD, lung function, pulmonary function, post bronchodilator FEV1, asthma) and “mood/brain-related” (i.e. brain region volumes, cortical surface area, depressed affect, neuroticism, Parkinson’s disease, white matter microstructure, smoking behaviour). Genes interacting with COPD-associated genes (level 1) within LSPPIN are regulated by eQTLs that have previously been associated with Alzheimer’s disease, atopic asthma and post bronchodilator FEV1 or FEV1/FVC ratio. Level 2-genes are only regulated by eQTLs previously associated with “mood/brain-related” (e.g. depressed affect, neuroticism, Parkinson’s disease) or “blood-related” (e.g. blood protein levels) traits. Genes within levels 3 and 4 are mostly associated with eQTLs enriched within “blood-related” traits.

### COPD shows associations with lung function, CAD, AD and brain region volumes

Four COPD-associated genes (i.e. *MSL1, MOCS2, NUPR1* and *SGF29*) were involved in the sulfur relay system and folate biosynthesis pathways that have previously been linked to COPD (Stepaniants et al. 2014; Kim et al. 2020; Shi et al. 2022). These four genes are associated with eQTLs linked to COPD and lung function (Fig. 4). Notably, *MSL1, MOCS2, NUPR1* and *SGF29* interact with three genes that are associated with post bronchodilator FEV1, atopic asthma and Alzheimer’s disease (Fig. 4A).

**Figure 4.**
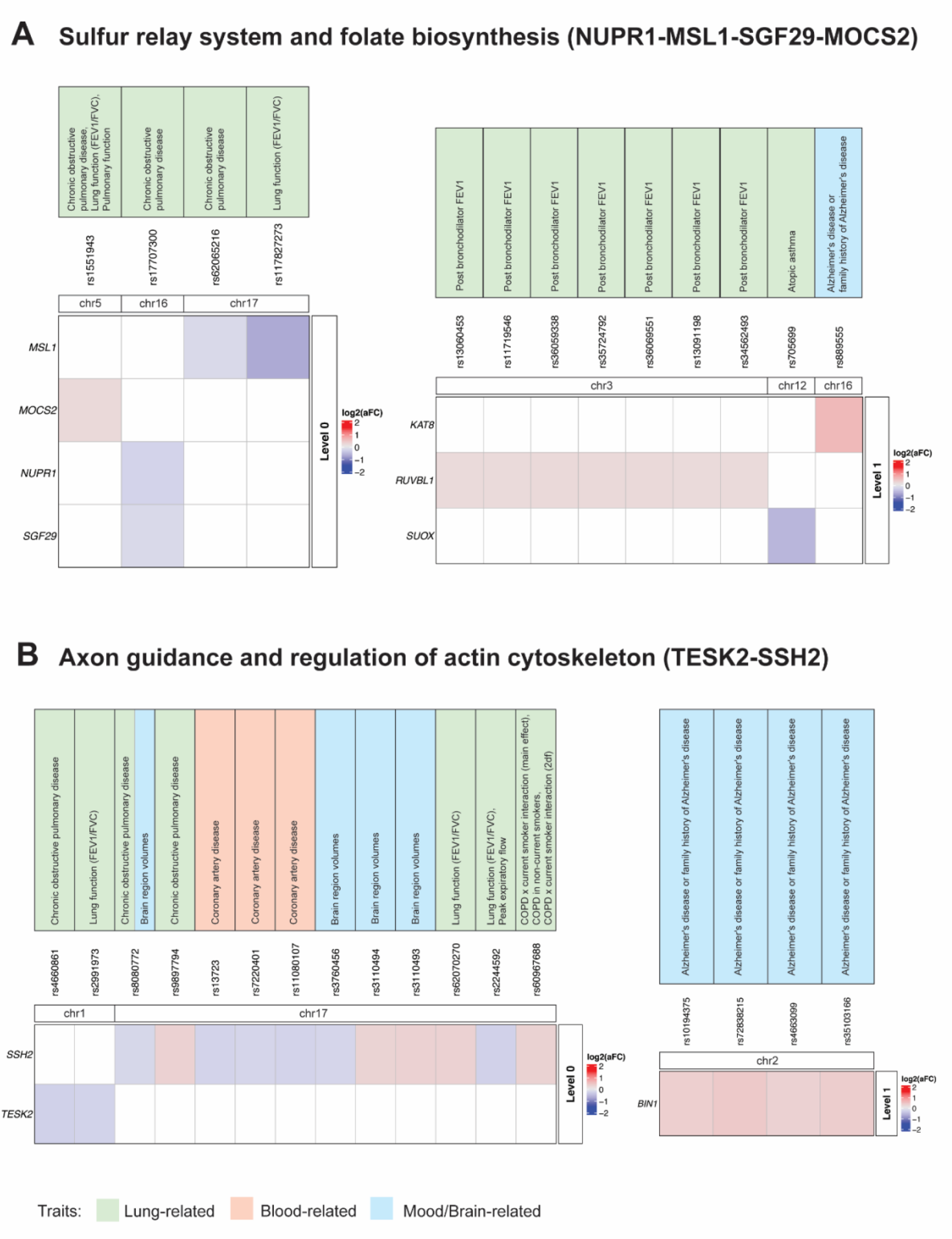
Trait-eQTL-gene associations within levels 0 and 1 for two COPD-associated biological pathways. (**A)** Within the “Sulfur relay system and folate biosynthesis” PPI cluster four genes (i.e. *MSL1, MOCS2, NUPR1* and *SGF29*) are associated with eQTLs linked to COPD and lung functioning. These four genes interact with three genes (“level 1”) that are associated with post bronchodilator FEV1, atopic asthma and Alzheimer’s disease. (**B)** *SSH2* and *TESK2* within the “Axon guidance and regulation of actin cytoskeleton” PPI cluster are associated with eQTLs linked to COPD, lung functioning, brain region volumes and CAD. Both *SSH2* and *TESK2* genes interact with *BIN1* gene (“level 1”), that is associated with Alzheimer’s disease.

Within the “Axon guidance and regulation of actin cytoskeleton” protein cluster, *SSH2* and *TESK2* are associated with eQTLs linked to COPD, lung functioning, brain region volumes and CAD (Fig. 4B). SSH2 and TESK2 proteins interact with the BIN1 protein (“level 1”), which has been associated with Alzheimer’s disease.

Reversing the analysis, using CAD-associated (n=804) and UD-associated (n=932) SNPs (*p*<5×10^−8^) confirmed COPD was co-occuring with CAD (“level 0”, Supplemental Fig. S5). By contrast, analysis of UD identified general lung function (FEV1/FVC), asthma and “mood/brain-related” (i.e. bipolar disorder, depression, autism spectrum disorder, schizophrenia) as being linked to UD-associated genes (Supplemental Fig. S6).

## Discussion

We integrated genes that are targeted by spatially constrained COPD-associated eQTLs with a lung-specific GRN, LSPPIN and all GWAS SNP-trait associations to identify traits that are co-occuring with COPD. The results of this integration provide insights into the regulatory mechanisms underlying these associations. We identified co-occuring traits that have been previously linked to COPD (e.g. lung function, asthma, depressed affect, CAD, AD, smoking behaviour, Parkinson’s disease (Burke and Wilkinson 2021; Carmona-Pírez et al. 2021; Xia et al. 2020; Martucci et al. 2021; Ställberg et al. 2018; Li et al. 2015; Cavaillès et al. 2013)) and those that have not (brain region volumes and white matter microstructure). We contend that the total eQTLs we identified, as impacting on COPD and its co-occuring traits, represent the ‘total’ genetic burden that contributes to an individual’s risk of developing COPD and its diverse clinical presentations.

The results of this study should be interpreted in view of its strengths and limitations. The main strength of this study is the integration of independent datasets: lung-specific 3D genome structure (Schmitt et al. 2016), common GWAS SNPs, genotypes and gene expression data (Aguet et al. 2019), and protein-protein interaction (Szklarczyk et al. 2019) data. Integrating these datasets enabled the identification of the impact of spatially constrained COPD-associated eQTLs on genes and biological pathways that link to co-occuring conditions. Indeed it is possible that the ‘total’ genetic burden we identified can be used to stratify COPD and it’s multiple clinical presentations, which have previously led to questions about the validity of classifying it as a single diagnostic category (Alabi et al. 2021; Corlateanu et al. 2020; Sakornsakolpat et al. 2019). However, this study also has several limitations. Firstly, this study was focused on the regulatory roles of common genetic variants (MAF ≥ 0.05) ignoring the impacts of other genetic (e.g. rare SNPs) and environmental factors, which will undoubtedly contribute to the risk of COPD and its co-occuring conditions. Secondly, we focused on the extended protein-protein interaction network within the lung gene regulatory network, as the lung represents the primary affected tissue in COPD. However, it is possible that genetic variation will impact on COPD risk through other tissues (e.g. blood) (Burke and Wilkinson 2021). Thirdly, the tools and datasets used in this study are potentially biased (e.g. lung-specific eQTL and Hi-C data were not obtained from identical samples). Furthermore, the identification of the co-occuring conditions was limited to the traits that have been studied and were present in the GWAS catalog. Finally, mapping to Ensembl gene identifiers potentially causes a loss of data specificity, since alternative splicing typically produces multiple transcripts and protein variants.

Co-occuring conditions are commonly associated with COPD and increase the risk of hospitalisation (Schnell et al. 2012). Our analysis of co-occuring conditions has identified risk variants and protein interactions that connect COPD with smoking (Hopkinson 2022), asthma (Maselli and Hanania 2018), coronary artery disease (CAD) (Xia et al. 2020), lung cancer (Parris et al. 2019), multiple sclerosis (Egesten et al. 2008), kidney failure (Trudzinski et al. 2019), unipolar depression (UD) (Li et al. 2019), Alzheimer’s disease (AD) (Wang et al. 2019), Parkinson’s disease (PD) (Li et al. 2015), personality traits (e.g. neuroticism) (Terracciano et al. 2017; Caille et al. 2021; Chetty et al. 2017). These conditions have been previously reported as being co-occuring conditions in COPD patients but the biological basis of the connections was unknown.

Brain region volumes, white matter microstructure, left-right brain asymmetry and cortical surface area (global PC1) were additional traits that we have identified as being co-occuring with COPD. Notably, patterns of brain structural alteration have been reported in COPD with different levels of pulmonary function impairment and cognitive deficits (Yin et al. 2019). This suggests that COPD patients may exhibit progressive structural impairments in both the grey and white matter, along with impaired levels of lung function. In addition to the trait associations, the results of our study also provide evidence for the putative genetic and biological connections between these traits.

Impaired lung function as measured by forced vital capacity (FVC) or forced expiratory volume in the first second (FEV1) has been previously associated with insulin resistance (Machado et al. 2018; Piazzolla et al. 2017; Sagun et al. 2015). Previous studies have also identified elevated levels of plasma transforming growth factor-β (TGF-β), an important regulator of lung and immune system development, in COPD patients compared to healthy controls (Mak et al. 2009; Verhamme et al. 2015). We identified COPD-associated genes that are involved in insulin resistance and immune pathways. In addition, we identified COPD-associated eQTLs that may target cholinergic pathway genes (e.g. *CHRM1, CHRM3*) implicated as important susceptibility loci for lung diseases (e.g. asthma, COPD) (Rajasekaran et al. 2019; Palmberg et al. 2018). The cholinergic pathway mediated by the parasympathetic neurotransmitter, acetylcholine, is a predominant neurogenic mechanism contributing to bronchoconstriction (Ward 2022). Notably, changes in the parasympathetic neuronal control of airway smooth muscle have been shown to increase bronchoconstriction in response to vagal stimulation, leading to airway hyperresponsiveness (Ward 2022). At the biological level, these findings emphasize the effects that COPD-associated SNPs may have on the regulation of genes within specific biological pathways (e.g. through creating an imbalance in the concentration of specific proteins), which, in turn, can be associated with an increased risk of disease.

In conclusion, we have integrated different levels of biological information (i.e. genes that are targeted by spatially constrained COPD-associated eQTLs, lung-specific GRN, LSPPIN and all GWAS SNP-trait associations) to identify target genes, associated with COPD-associated eQTLs, that interact to connect COPD to co-occuring conditions. Collectively, these results provide multiple new avenues for investigation of the underlying biology and diverse clinical presentations of COPD, and suggest potential therapeutic COPD markers necessary for the follow-up patient stratification.

## Methods

### Hi-C data processing

Hi-C chromatin interaction libraries specific to primary lung cells (Supplemental Table S1) were downloaded from GEO database (https://www.ncbi.nlm.nih.gov/geo/, accessions: GSM2322544 and GSM2322545) and analyzed using the Juicer pipeline (v1.5) (Durand et al. 2016) to generate Hi-C libraries. The pipeline included BWA (v0.7.15) alignment of paired-end reads onto the hg38 (GRCh38; release 75) reference genome, merging paired-end read alignments and removing chimeric, unmapped and duplicated reads. We refer to the remaining read pairs as “contacts”. Only Hi-C libraries that contain >90% alignable unique read pairs, and >50% unique contacts (<40% duplication rate) within the total sequenced read pairs were included in the analysis. Files containing cleaned Hi-C contacts (*i*.*e*. *_merged_nodups.txt files) were processed to obtain Hi-C chromatin interaction libraries in the following format: read name, str1, chr1, pos1, frag1 mapq1, str2, chr2, pos2, frag2, mapq2 (str = strand, chr = chromosome, pos = position, frag = restriction site fragment, mapq = mapping quality score; 1 and 2 correspond to read ends in a pair). Reads where both ends had a mapq ≥30 were included in the final library. Hi-C chromatin interactions represent all captured pairs of interacting restriction fragments in the genome and were used by CoDeS3D to identify putative regulatory interactions between SNPs and genes.

### Identification of SNPs associated with COPD, CAD and UD

Single-nucleotide polymorphisms (SNPs) associated (*p* < 5×10^−8^) with COPD (n=263), CAD (n=804) and UD (n=932) were downloaded from the GWAS Catalog (www.ebi.ac.uk/gwas/; 09/06/2021 and 11/04/2022; Supplemental Table S2).

### Identification of spatial regulatory interactions

The CoDeS3D (Fadason et al. 2018) pipeline was used to identify genes that spatially interact with putative regulatory regions tagged by SNPs (Fig. 1A). Briefly, the human genome build hg38 (GRCh38; release 75) was fragmented *in silico* at HindIII sites (A^AGCTT), the restriction enzyme that was used in the preparation of the lung-specific Hi-C libraries (Schmitt et al. 2016). Disease associated SNP rsID numbers were cross-checked with the GTEx v8 lung eQTL database (GTEx Consortium 2020) and restriction fragments that were tagged by the COPD associated SNPs were identified. Using lung-specific Hi-C libraries (Supplemental Table S1), CoDeS3D identified the restriction fragments that were captured interacting with the SNP-tagged restriction HindIII fragments. Interacting fragments that overlapped annotated genes (GENCODE transcript model version 26) were identified. The resulting SNP-gene pairs were used to query the GTEx v8 lung eQTL database (GTEx Consortium 2020) to identify cis- and trans-acting eQTLs (*i*.*e*. genes, whose expression levels are associated with the SNP identity). Finally, significant lung-specific eQTL-gene interactions were identified using the Benjamini-Hochberg (BH) FDR correction to adjust the eQTL *p* values (FDR<0.05).

CoDeS3D was used to build a lung-specific gene regulatory network (GRN, Supplemental Fig. S2). All SNPs (MAF ≥ 0.05, n=43,174,097) present within GTEx lung-specific eQTL database (GTEx Consortium 2020) were used to identify all significant (BH, FDR < 0.05) spatially constrained lung-specific eQTL-gene interactions (the lung-specific GRN). Multiple correction testing was performed across all interactions within individual chromosomes.

The lung-specific GRN was mined using the COPD-associated SNPs (n=263; *p* < 5×10^−8^; GWAS Catalog; 09/06/2021) to identify all COPD-associated significant (BH, FDR<0.05) lung-specific eQTL-gene interactions (COPD-associated GRN) (Supplemental Table S3). This was repeated for the CAD-associated (n=651; *p* < 5×10^−8^; GWAS Catalog; 11/05/2022) and UD-associated (n=152; *p* < 5×10^−8^; GWAS Catalog; 11/05/2022) SNPs (CAD- and UD-associated GRNs, respectively; Supplemental Tables S4 & S5).

### Functional annotation of eQTL SNPs associated with COPD

The COPD-associated eQTLs were annotated using the wANNOVAR tool (Chang and Wang 2012) (http://wannovar.wglab.org/, 09/06/2021) to obtain information about the locus they tagged (Supplemental Table S2). All genomic positions and SNP annotations were obtained for human genome reference build hg38 (GRCh38) release 75.

### Construction of lung-specific protein-protein interaction network

The STRING (Szklarczyk et al. 2019) protein-protein interaction (PPI) database (version 11.5, https://string-db.org/, 15/03/2022) was downloaded and queried (STRING API) to identify potential protein-protein interactions (combined score ≥0.7). A lung-specific PPI network (LSPPIN) was constructed by filtering the STRING PPI network for the proteins encoded by the genes that were affected by eQTLs within the lung-specific GRN (Supplemental Fig. S3). Ensembl protein identifiers were mapped to Ensembl gene identifiers using EnsDb.Hsapiens.v86 R package. The LSPPIN represents a subnetwork of the entire STRING PPI network, in which a protein/node is only present if the encoding gene is associated with a spatially constrained eQTL within lung tissue. The size of each node depends on the protein expression levels (no missing values, TPM >0.1 and ≥6 reads in a minimum of 20% of tested samples) within the GTEx v8 lung database (GTEx Consortium 2020). The resulting LSPPIN contained 210,192 PPIs between 10,188 unique proteins.

### Construction of COPD-associated LSPPIN

To build the COPD-specific LSPPIN, only interactions between genes targeted by COPD-associated eQTLs were extracted from the LSPPIN (Supplemental Table S6). Louvain clustering (Blondel et al. 2008) was applied to identify clusters of functionally related proteins within the COPD-associated LSPPIN.

### Identification of potential co-occuring conditions

The Multimorbid3D pipeline was used to identify traits that were co-occuring with COPD (Fig. 1B) (Zaied et al. 2022). Briefly, ‘all’ eQTLs that were associated with the proteins encoded by the genes targeted by the COPD-eQTLs (level 0) were identified within the LSPPIN. These steps were repeated for the genes that encoded proteins that were identified as interacting with the COPD-associated proteins (level 1), genes encoding the proteins interacting with “level 1” proteins (level 2), “level 2” genes (level 3), and “level 3” genes (level 4) within the LSPPIN. The GWAS Catalog (www.ebi.ac.uk/gwas/, v1.0.2, 30/03/2022) was queried to identify traits that were enriched for the eQTLs from each level. The hypergeometric distribution test was used to identify significant enrichment of traits at each level within the GWAS catalog traits. Finally, significantly enriched traits were identified using the BH FDR correction to adjust the *p* values (FDR ≤ 0.05; Supplemental Table S7). The same pipeline was also used to identify the co-occuring conditions for CAD and UD (Supplemental Table S7).

### Gene Ontology enrichment and pathway analyses

Gene Ontology (GO) enrichment and pathway analyses were performed using the g:GOSt module of the g:Profiler tool (Raudvere et al. 2019) (Supplemental Tables S8 & S9). Enrichment was tested for within the biological process, molecular function and cellular component GO terms. All annotated human genes were chosen as the statistical domain scope. The significance level was determined using the BH algorithm (FDR < 0.05). The Kyoto Encyclopedia of Genes and Genomes (KEGG) database (Kanehisa and Goto 2000) was used to identify impacted biological pathways.

### Bootstrapping analysis

Bootstrapping (n=1000 iterations) was performed to test the specificity of the identified genes, pathways and traits to COPD. The *p*-value was calculated as the number of the occurrences of COPD-associated genes in the iterations divided by 1000.

### Data access

Data access was approved by the dbGaP (https://www.ncbi.nlm.nih.gov/gap/) Data Access Committee(s) for total RNA-seq and WGS datasets across GTEx v8 tissues (project #22937: “Untangling the genetics of disease multimorbidity”, accession: phs000424.v8.p2) (Aguet et al. 2019) (Supplemental Table S1). Data analysis and visualisation were performed in Python (version 3.6.9) using miniconda (version 4.8.4), and R (version 4.0.2) through RStudio (version 1.2.5033). All datasets and software used in the analysis are listed in Supplemental Table S1. All findings, scripts and a reproducibility report are available https://github.com/Genome3d/genetic_regulation_in_COPD.

Juicer is available at https://github.com/aidenlab/juicer.

CoDeS3D is available at https://github.com/Genome3d/codes3d-v2.

Multimorbid3D is available at https://github.com/Genome3d/multimorbid3D.

## Acknowledgements

The authors would like to thank the Genomics and Systems Biology Group (Liggins Institute, University of Auckland) for useful discussions. The authors wish to acknowledge the use of New Zealand eScience Infrastructure (NeSI, https://www.nesi.org.nz) high performance computing facilities, consulting support and/or training services as part of this research. New Zealand’s national facilities are provided by NeSI and funded jointly by NeSI’s collaborator institutions and through the Ministry of Business, Innovation & Employment’s Research Infrastructure programme. This work was funded by the Dines Family Charitable Trust. The Genotype-Tissue Expression (GTEx) Project was supported by the Common Fund of the Office of the Director of the National Institutes of Health, and by NCI, NHGRI, NHLBI, NIDA, NIMH, and NINDS.

## Author contributions

E.G. performed data analyses, created visualizations and wrote the manuscript. T.F. contributed to data interpretation and manuscript revision. R.K.J. contributed to data analysis and manuscript revision. H.K., J.J., K.B., M.T. contributed to manuscript revision. J.M.O. directed the study, contributed to data interpretation and co-wrote the manuscript.

