## Supplemental figures for "*De novo* discovery of traits co-occurring with chronic obstructive pulmonary disease"

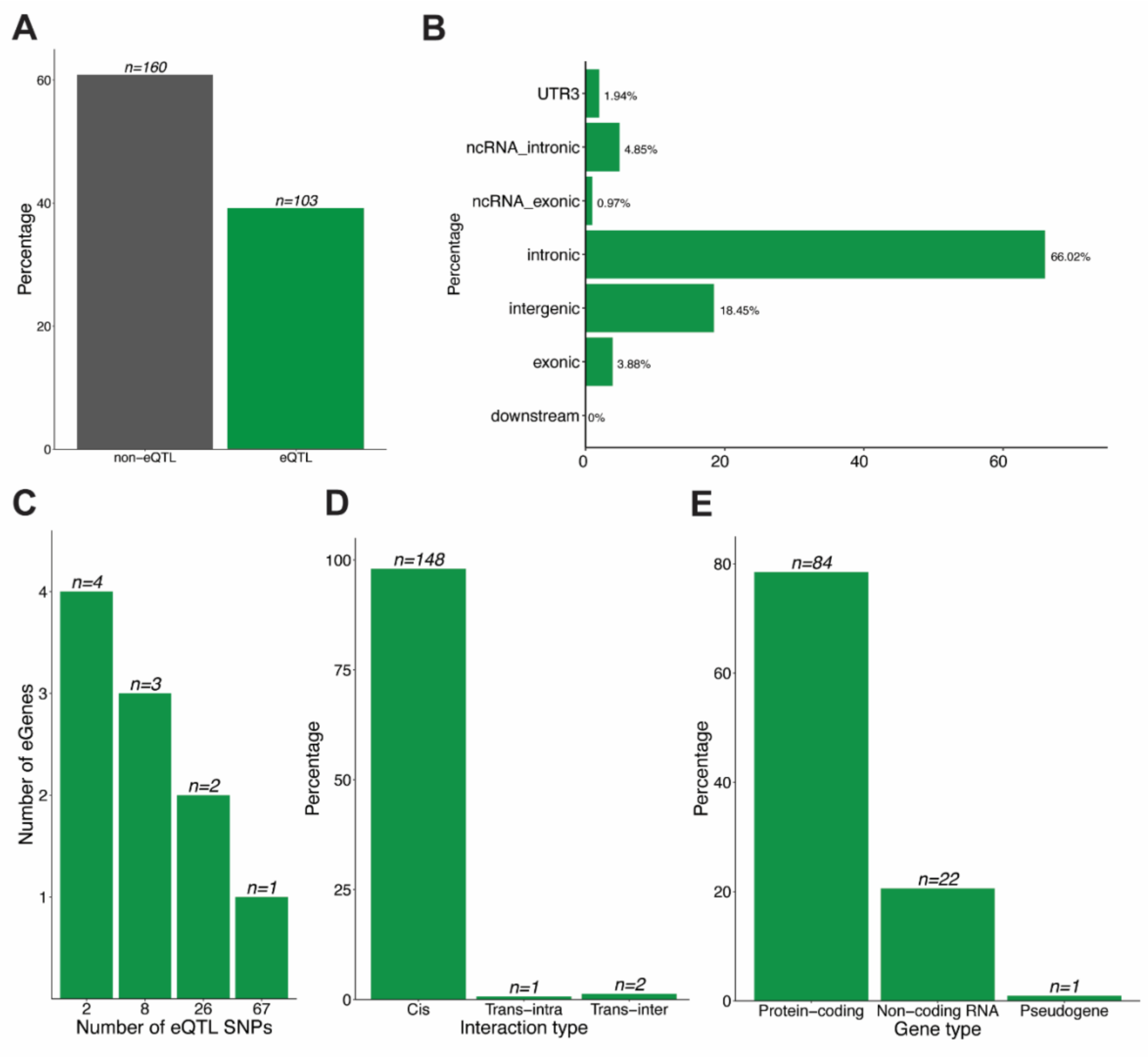


**Supplemental figure S1.** Characteristics of COPD-associated lung-specific GRN. (**A)** Of 263 COPD-associated GWAS SNPs, 103 SNPs are involved in spatially constrained eQTL-gene interactions in the lung. (**B)** Approximately 96% of the COPD-associated eQTLs are located within non-coding genomic regions (Supplemental Table S2). (**C)** The majority of COPD-eQTLs (n=67) associate with the transcript levels of one gene. Two eQTLs (i.e. rs2277027 and rs9435731) control four genes (i.e. *ADAM19*, *CTB-109A12.1*, *CTB-47B11.3*, *CYFIP2* and *ATP13A2*, *CROCC*, *MFAP2*, *RP1-37C10.3*, correspondingly). (**D)** The majority of the eQTL-gene interactions (n=148) are cis-acting. (**E)** The majority of COPD-eQTLs are associated with changes in transcript levels of protein-coding genes (n=84) within the lung.

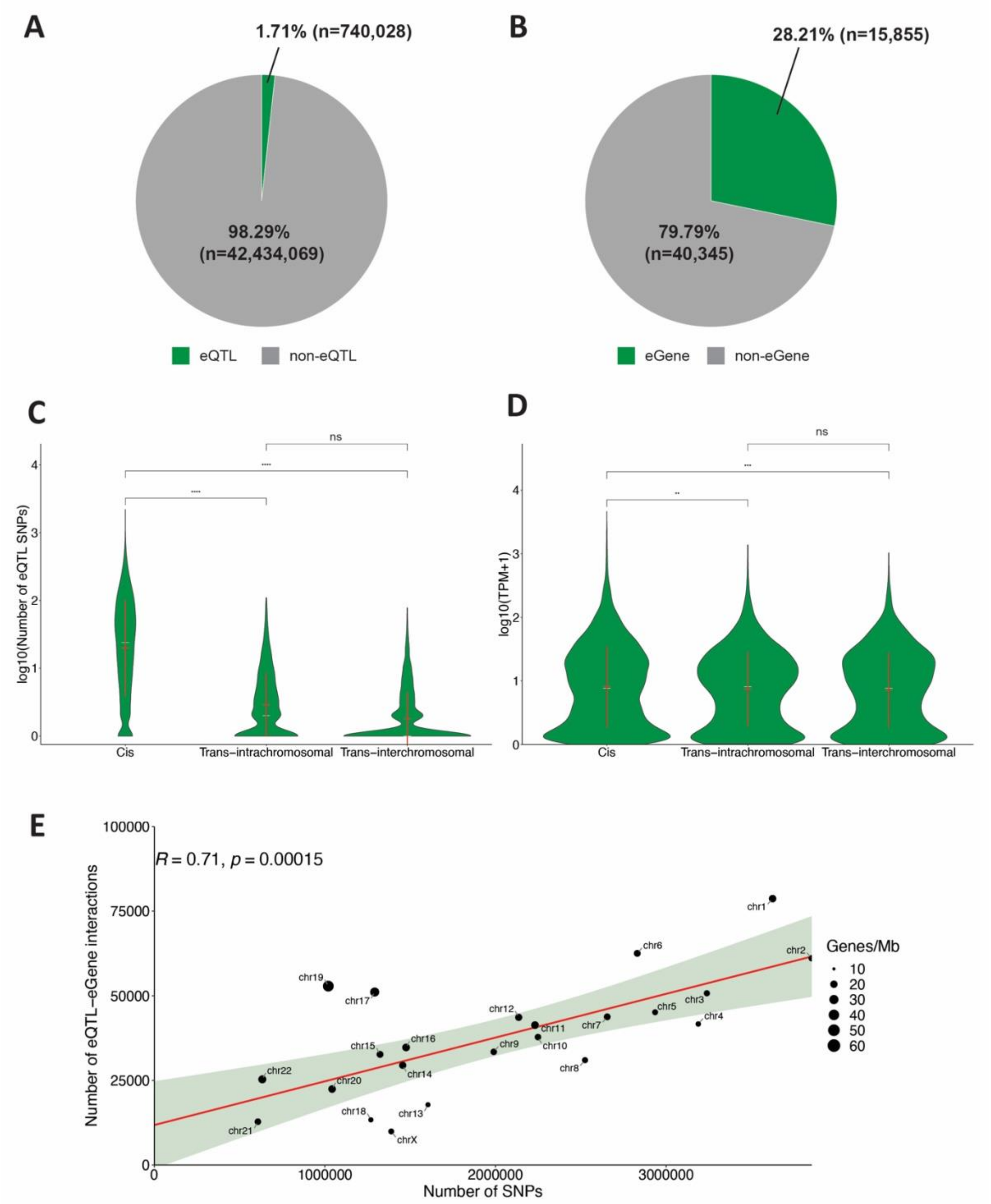


**Supplemental figure S2.** Characteristics of the lung-specific gene regulatory network (GRN). (**A)** Out of 43,174,097 genetic variants present within the GTEx database, only 1.71% (n=740,028) are involved in 873,133 spatially constrained eQTL-gene regulatory interactions in the lung tissue. (**B)** Of the 56,200 genes present in the GTEx database, 28.21% (n=15,855) are involved in spatially constrained eQTL-gene regulatory interactions within the lung tissue. (**C**) Of the spatially constrained eQTL-gene regulatory interactions, 850,093 are cis (between 720,083 eQTLs and 15,044 genes), 16,414 are trans-intrachromosomal (between 16,376 eQTLs and 2,743 genes) and 6,626 are trans-interchromosomal (between 6,469 eQTLs and 2,024 genes). There are significantly more cis interactions than trans-interactions (t test, $p \leq 0.0001$). Mean (red line), median (white line) and standard deviation of the number of spatially constrained eQTLs per gene grouped by eQTL interaction type (i.e. cis, trans-intra, trans-inter). (**D)** Genes associated with cis-acting eQTLs have higher expression levels than those associated with trans- interactions (t test, $p \leq 0.01$ and $p \leq 0.001$, respectively). Mean (red line), median (white line) and standard deviation of gene transcript levels grouped by eQTL interaction type. **(E)** Correlation analysis between the number of spatially constrained eQTL-gene interactions from the lung-specific GRN and the number of all common (MAF $\geq$ 0.05) GTEx genetic variants identifies chromosomes X, 18, 13, 8 and 4 as having fewer eQTLs than expected. By contrast, chromosomes 19, 17, 6 and 1 tend to have more eQTLs than expected in the lung tissue, with chromosomes 6 and 17 exhibiting the greatest deviation from the prediction curve. t test: ns, not significant ($p > 0.05$), ** significant ($p \leq 0.01$),*** significant ($p \leq 0.001$),**** significant ($p \leq 0.0001$).

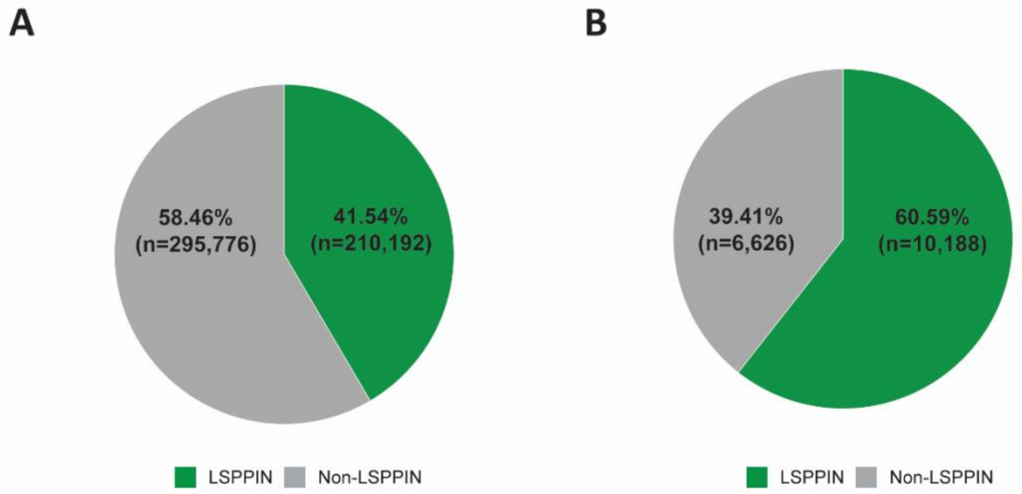


**Supplemental figure S3.** Characteristics of lung-specific protein-protein interaction network (LSPPIN). The proteins encoded by all genes (n=15,855) within the whole lung-specific GRN were mapped to the curated STRING human protein-protein interaction database (combined score $\geq$ 0.7). (**A)** Of 505,968 STRING human PPIs, 41.54% (n=210,192) are present within the lung-specific GRN. (**B)** Of 16,814 human unique proteins in the STRING database, 60.59% (n=10,188) are present in the lung tissue.

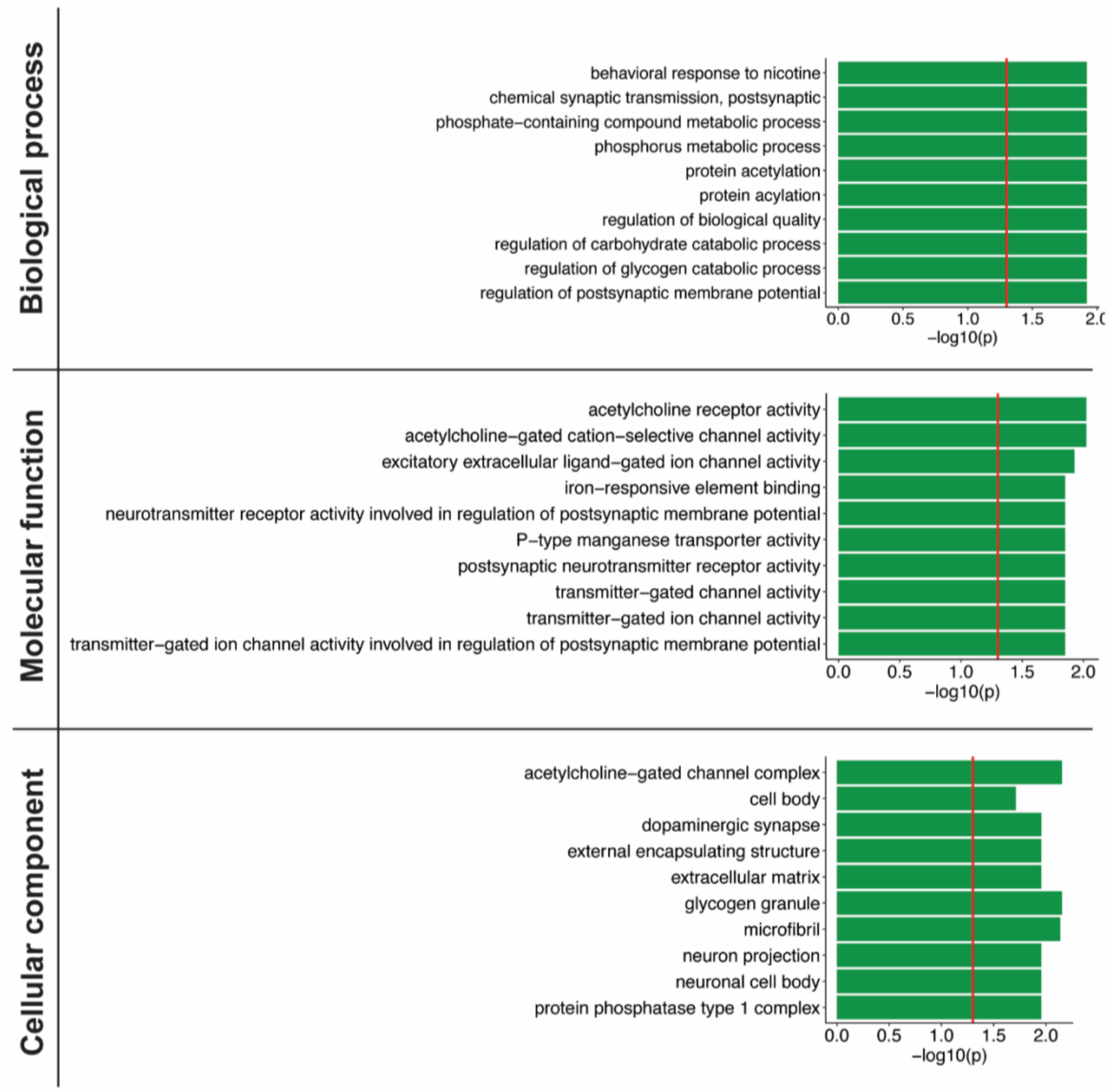


**Supplemental figure S4.** Top 10 biological processes, molecular functions and cellular components for genes that are targeted by spatially constrained COPD-associated eQTLs within the lung. Gene ontology (GO) enrichment analysis was performed using g:Profiler. The threshold for significance (red line) is $p<0.05$.

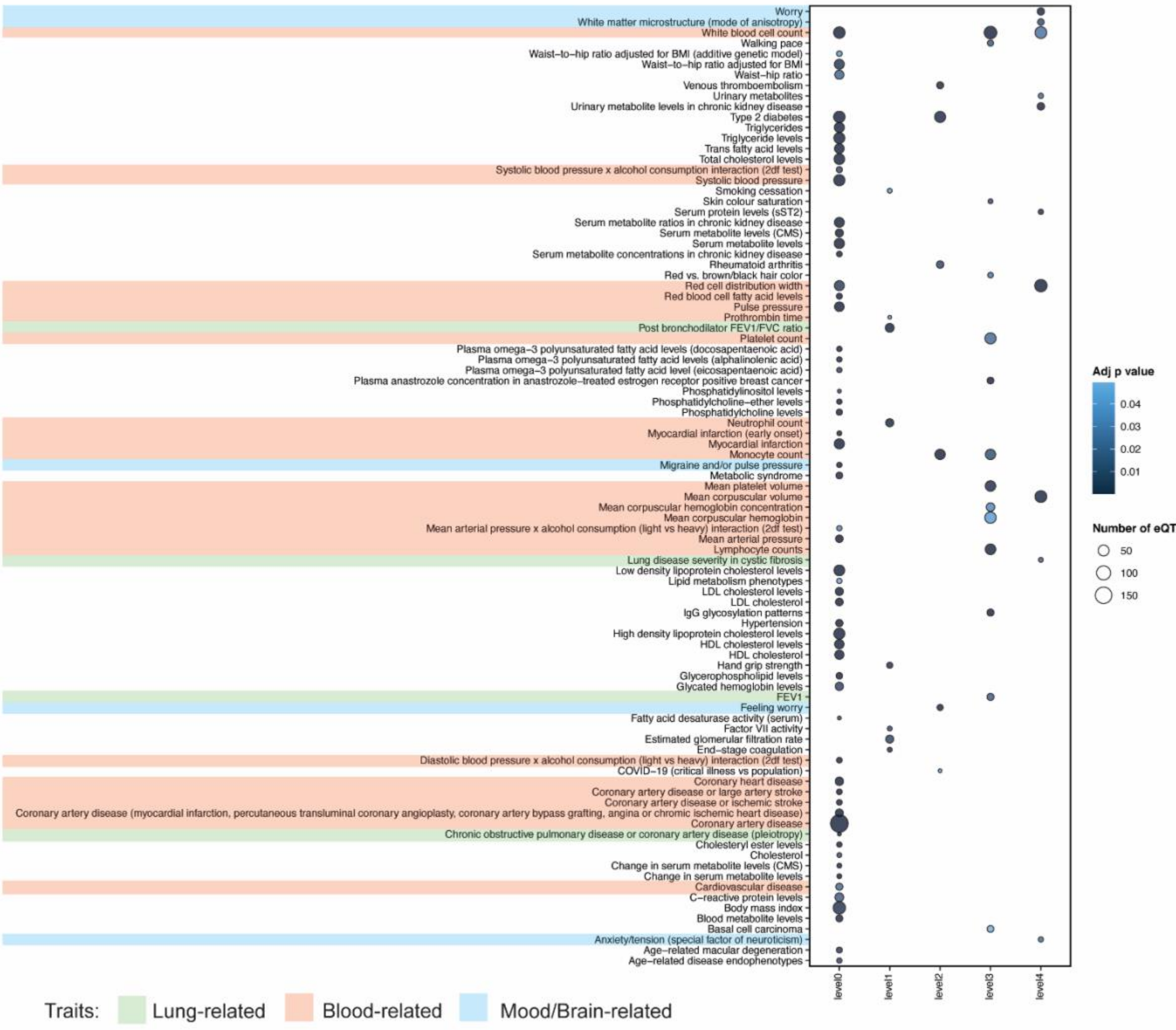


**Supplemental figure S5**. Co-occuring conditions that are putatively associated with CAD. We identified 58 GWAS traits that are enriched (FDR≤0.05) for eQTLs associated with CAD-eQTL targeted genes (level 0) (Supplemental Table S6). Among these co-occuring traits are "blood- and heart-related" (i.e. CAD, coronary heart and cardiovascular diseases). COPD is also found to be associated with CAD-eQTL targeted genes (level 0, "Chronic obstructive pulmonary disease or coronary disease (pleiotropy)").

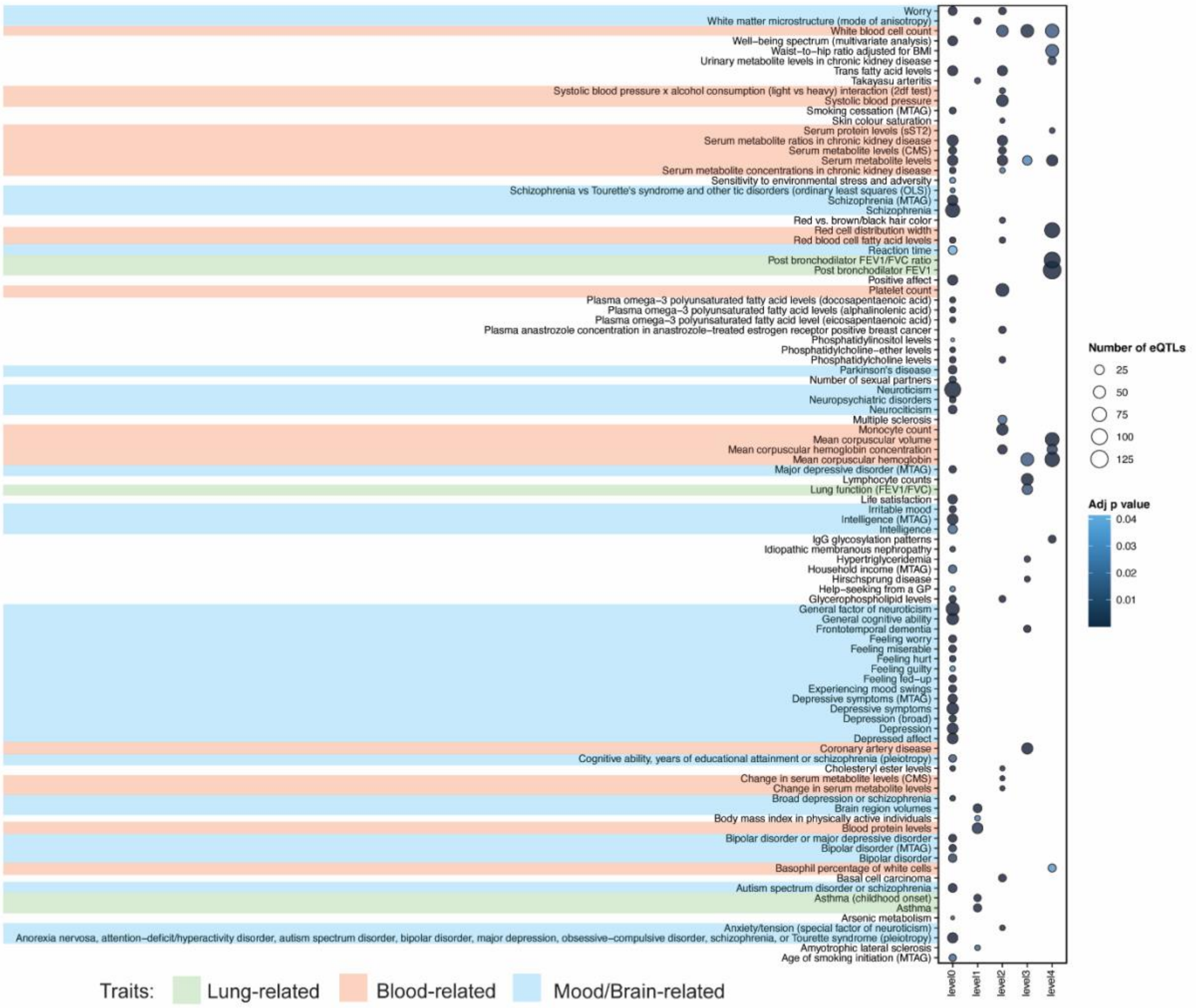


**Supplemental figure S6**. Co-occuring conditions that are putatively associated with UD. We identified 58 GWAS traits that are enriched (FDR≤0.05) for eQTLs associated with UD-eQTL targeted genes (level 0) (Supplemental Table S6). As expected, the majority of the identified co-occuring traits on level 0 are "mood/brain-related" (i.e. Bipolar disorder, Depression, Autism spectrum disorder or schizophrenia). However, a few "lung-related" are also found to be associated with UD-eQTL targeted genes (level 0, Lung function (FEV1/FVC), Asthma).
